# vascr: An R-based toolkit for rapid and robust analysis of cellular impedance sensing data

**DOI:** 10.64898/2025.12.17.695026

**Authors:** James J. W. Hucklesby, Charles P. Unsworth, E. Scott Graham, Catherine E. Angel

## Abstract

**Purpose:** Electrical impedance sensing offers a powerful, non-invasive method for real-time monitoring of cellular behaviour, with applications spanning immunology, cancer biology and drug discovery. Adoption of impedance-sensing approaches has been accelerated by the commercial availability of the ECIS, xCELLigence, ScioSpec and cellZscope platforms; however, the complexity and volume of impedance datasets produced has limited rigorous analysis.

**Methods:** vascr, an open-source R-based toolkit designed to streamline the processing of impedance sensing datasets was created to facilitate rapid and simplified handling of cellular impedance sensing data.

**Results:** vascr can import multi-frequency data from multiple instrument manufacturers and includes robust analysis tools for resampling, outlier detection, and statistical testing using ANOVA and cross-correlation.

**Conclusions:** Using the vascr workflow, impedance sensing data can be rapidly and robustly interpreted allowing for real-time investigations of the effects of biological agents on cellular processes such as endothelial barrier integrity.

## 1. Introduction

Electrical impedance measurements provide a non-invasive method for measuring cell function in real-time. Impedance sensing measurements are taken by culturing cells on conductive electrodes, allowing measurement of quantitative impedance spectra [1–6]. As the currents used for these measurements are small, they are undetectable to the cells, repeated measurements can be taken rapidly without disturbing the culture, providing a high temporal resolution over an extended period [2, 5–8]; far exceeding the resolution and duration of Transwell tracer, EVOM TEER and live cell or surgical imaging techniques [6, 9–11]. Furthermore, the measurement electronics can be multiplexed, allowing for up to 96 wells to be monitored simultaneously [6]. Hence, electrical impedance experiments are ideally suited to monitoring large numbers of treatments with biological responses which change over time [12, 13]. These have been applied to a wide variety of applications, including monitoring cellular differentiation [14], cancer growth [15, 16], wounding [17, 18], renal research [19], metastatic invasion [20, 21], blood-brain barrier dysfunction[6, 20, 22, 23] cell migration[4, 5, 24] and drug responses [13]. Furthermore, the emergence of three leading commercial options; ECIS, xCELLigence and cellZscope has made the technology accessible to laboratories without extensive electrical engineering experience [6].

Once the impedance measurements have been collected, various biological characteristics of the cells can be inferred. In its simplest form, electrical parameters measured at specific frequencies can be linked to biologically significant characteristics. Measurements of overall barrier integrity, like those generated by traditional TEER assays, can be mimicked by measuring resistance at 4000 Hz [8]. Hence, this simple and robust measurement has been widely used to assess the integrity of cellular monolayers in studies of inflammation, cancer, immunology and cardiology [6, 16–18, 20, 21, 23, 25]. Additionally, electrical capacitance provides a proxy for cellular proliferation and has been used to measure the proliferation of cancer cells and to monitor cell viability during viral infection [1, 8, 15, 25, 26]. Whilst measuring single frequencies is straightforward, this approach lacks the sensitivity to detect biological processes which affect a wide range of frequencies to a small extent, such as cell-cell interactions or basolateral adhesion [6, 10, 25].

More nuanced biological insights can be made by fitting a mathematical model of an equivalent circuit to impedance values measured at a range of frequencies [6, 8]. One highly utilised model is that described by Giaever and Keese, which assesses the properties of endothelial monolayers [1, 8]. When applying this model, values representing cell-cell interactions (Rb), membrane capacitance (Cm), and basolateral adhesion (α) can be individually determined. Each of these modelled values is driven by different cellular processes, increasing the specificity of the results [6, 25]. This approach has been used in a wide variety of studies to monitor multiple barriers, including those within the brain[25], lungs [27], intestine [17] and retina [28]. Within our group, we have utilised this technology to monitor the integrity of brain endothelial monolayers in the presence of cytokines, proteases and melanoma cells [6, 11, 20–22].

However, impedance sensing is ultimately not a direct measurement of biological function but a measure of the electrical properties of cells. Hence, although measuring electrical properties allows for impedance sensing’s characteristic accuracy, throughput and temporal resolution, this electron flow is not restricted to any single biological process [4, 5, 10, 29]. In contrast, techniques such as immunocytochemistry or Transwell tracer assays are far more specific, as they measure fluorophores covalently linked to molecules involved specifically in a particular biological process [10]. Therefore, the gold standard approach remains to trade off the relative strengths of both approaches, utilising impedance sensing to determine when and where the biology is changing and confirming results using direct labelling techniques [6, 10].

Whilst impedance sensing is technically complex, commercially available platforms have gained widespread use and increased the accessibility of impedance sensing [6, 30]. However, processing the impedance sensing data remains challenging. Typical ECIS measurements consist of many parameters, encompassing multiple electrical measurements, frequencies and wells. As these measurements are repeated every few minutes for several days, experiments usually span several hundred thousand datapoints[6]. Whilst these datasets can be processed manually, this approach is inherently time-consuming, prone to errors and hard to reproducibly replicate [31, 32]. Whilst commercially available instruments come with software to carry out basic analysis, the software is complex and differs substantially between manufacturers. Furthermore, this software often lacks the capability to combine multiple experiments and track experimental conditions, a requisite for conducting formal statistical analysis [12, 33, 34]. Hence, many investigators only utilise a small subset of the recorded impedance sensing measurements, present only a single representative experiment and forgo formal statistical testing [11, 35, 36]. When utilised, formal statistical analysis could enhance the interpretation of time series datasets by providing objective confirmation of the conclusions [37].

We have developed vascr, an R-based toolkit to rapidly and robustly analyse large impedance sensing datasets. In this paper, we will demonstrate how to use this toolkit to statistically assess real-time multi-frequency ECIS datasets. Whilst ECIS is used for the example presented in this paper, vascr will also analyse xCELLigence, ScioSpec and cellZscope data. Vascr can also perform two statistical tests: two-way ANOVA and cross-correlation. Two-way ANVOA analysis followed by Tukey’s HSD test provides a powerful method for objectively comparing the significance of differences between various treatments [38]. Hence, ANOVA has been applied to compare cellular impedance values at a single time point [6, 21, 39, 40]. Secondly, experimenters may wish to compare the temporal response between treatments over an extended period. Whilst this is commonly achieved by holistically comparing the shapes of two impedance sensing curves, differences are rarely quantified [6, 29, 41]. However, the similarity of the shapes between two temporal response curves throughout their entire duration can be calculated using cross-correlation analysis [6, 42]. Cross-correlation generates quantitative results, which can be utilised for formal statistical analysis using t-tests [43,44]. However, combined cross-correlation and ANOVA calculations are involved, and automated tools tailored for impedance sensing data remain underdeveloped.

In this paper, we present vascr, an integrated software solution for processing impedance-sensing data. vascr is built on the popular open-source R and Tidyverse packages, which have been used extensively to analyze large datasets [43, 44]. We demonstrate that vascr can import, graph and run statistical testing of impedance sensing data either directly through R, RStudio or via a web interface, allowing for rapid and robust analysis of impedance sensing data.

## 2. Methods

The example data used in this paper is a re-analysis of that presented in our 2021 publication [6]. A summary of how the data was collected is presented below.

### 2.1. Culture of brain microvascular endothelial cells

Human cerebral microvascular endothelial cells (hCMVECs) were purchased from Applied Biological Materials Inc (T0259) and used between passages 11 to 16. Cells were maintained in complete M199 medium (Life Technologies, 11150-059) supplemented with 10% FBS (Moorgate), 1 μg/mL hydrocortisone (Sigma-Aldrich, H0888), 3 ng/mL hFGF (Peprotech, AF-100-18B), 1 ng/mL hEGF (Peprotech, AF-100-15), 10 μg/mL heparin (Sigma-Aldrich, H3393-50KU), 2 mM GlutaMAX (Invitrogen, 35050061) and 80 μM dibutyryl-cAMP (Sigma-Aldrich, D0627) and stored at −80°C. Media not immediately required were stored at 4°C for no more than 14 days. Throughout all cultures and experiments, cells were maintained at 37 °C, with 5% CO2 and 100% humidity. All cell handling was performed under sterile conditions in a Class II hood.

### 2.2. Passage of cell lines

Cell lines were maintained and treated as previously described [6, 45]. Culture vessels were coated with 1 μg/cm² collagen I (Invitrogen, C3867) dissolved in 0.02 M acetic acid (Merck, 1.00063.2500) for 1 h at room temperature before being washed thrice with Type I water. T75 flasks containing adherent cells were washed twice with 4 ml pre-warmed PBS without calcium and magnesium (Gibco, 18912014) before being incubated with 4 ml pre-warmed TrypLE (Life Technologies, 12604021) for 5 min at 37°C. Cells were then visually inspected and tapped vigorously to ensure >90% detachment. 4 ml of complete media was added, and cells were triturated thrice. A 1ml aliquot of this suspension was added to a new pre-coated T75 flask with 12ml of media and returned to the incubator. Cells were passaged every 4 days. Routine mycoplasma testing was conducted using an eMyco+ PCR mycoplasma testing kit (LiliF Diagnostics, 25237).

### 2.3. Measuring hCMVEC monolayer properties using impedance sensing

96W20idf plates (Applied BioPhysics) were treated with 10 mM L-cysteine (Sigma-Aldrich, C7352) for 15 min. Wells were then coated with collagen, as described in the previous section.

hCMVEC cells, removed from the plate as previously described for maintenance, were pelleted at 100× g for 5 min, resuspended in 1 ml of complete media and enumerated using Trypan Blue and a haemocytometer. Unless otherwise stated, cells were seeded at a density of 62,500 cells/cm² in a volume of 150µl using a multi-channel pipette. After seeding, the plate was immediately installed into the ECIS 96 well array station housed within a cell culture incubator, and continuous measurements were initiated in multi-frequency mode using the default frequency spectra.

48 hours after seeding, 5x stocks of cytokine treatments or appropriate vehicle were prepared. ECIS readings were paused to introduce the treatment, and the plate was transferred to a polystyrene sheet within a class II hood. 44ul of 5x stock was gently introduced to the centre of each well. The plate was then returned to the ECIS 96 well array station, and measurements resumed for a further 48 hours.

### 2.4. Code availability

vascr is available from the CRAN repository (https://cran.r-project.org/web/packages/vascr) whilst the source code is available on GitHub (https://github.com/JamesHucklesby/vascr). Both repositories also contain full manuals, including worked examples of function usage. The web interface is accessible at http://vascr.huc.nz. Analysis was conducted using R (V 4.4.2) within RStudio (V 2024.12.0) [44, 46].

Whilst vascr provides a unified environment for impedance sensing, it depends on a range of packages to perform the underlying computation and plotting. Data was imported, cleaned and reshaped using rlang, dplyr, stringr, tidyr and utils packages [44, 47–50]. xCELLigence databases were imported with ODBC and DBI [51, 52]. Computationally expensive functions were accelerated using the foreach, furrr, doFuture and memoise packages [53–56]. Plots were generated using ggplot2, ggnewscale, ggpubr, ggrepell and ggtext packages [57–60]. Statistical comparisons were made using the multcomp, rstatx, stats and nlme packages [50, 61–63]. Progress updates and notifications were provided by cli and progressr [64, 65]. The web UI was implemented using Shiny, DT, bslib and shinyjs packages[66–69]. Data export was carried out by writexl [70]. The package is provided with full unit testing, which was implemented using testthat and vdiffr [71, 72].

## 3. Results and Discussion

### 3.1. vascr_import can import data from various instruments

Firstly, the vascr_import function was used to import data into a tibble. Although the ECIS, cellZscope and xCELLigence instruments store data in differing proprietary formats, vascr_import was able to import data from all three instruments into a consistently formatted tibble. This tibble contains all measurements and associated metadata collected by the instrument (Table 1). This includes both the raw impedance and phase measurements captured by the instrumentation, calculated resistance, reactance, and capacitance values and any modelled Cm, Rb and α values. As this data is in a tidy format, it can be conveniently piped into downstream R analyses to generate plots and conduct statistical analysis [43].

**Table 1.**
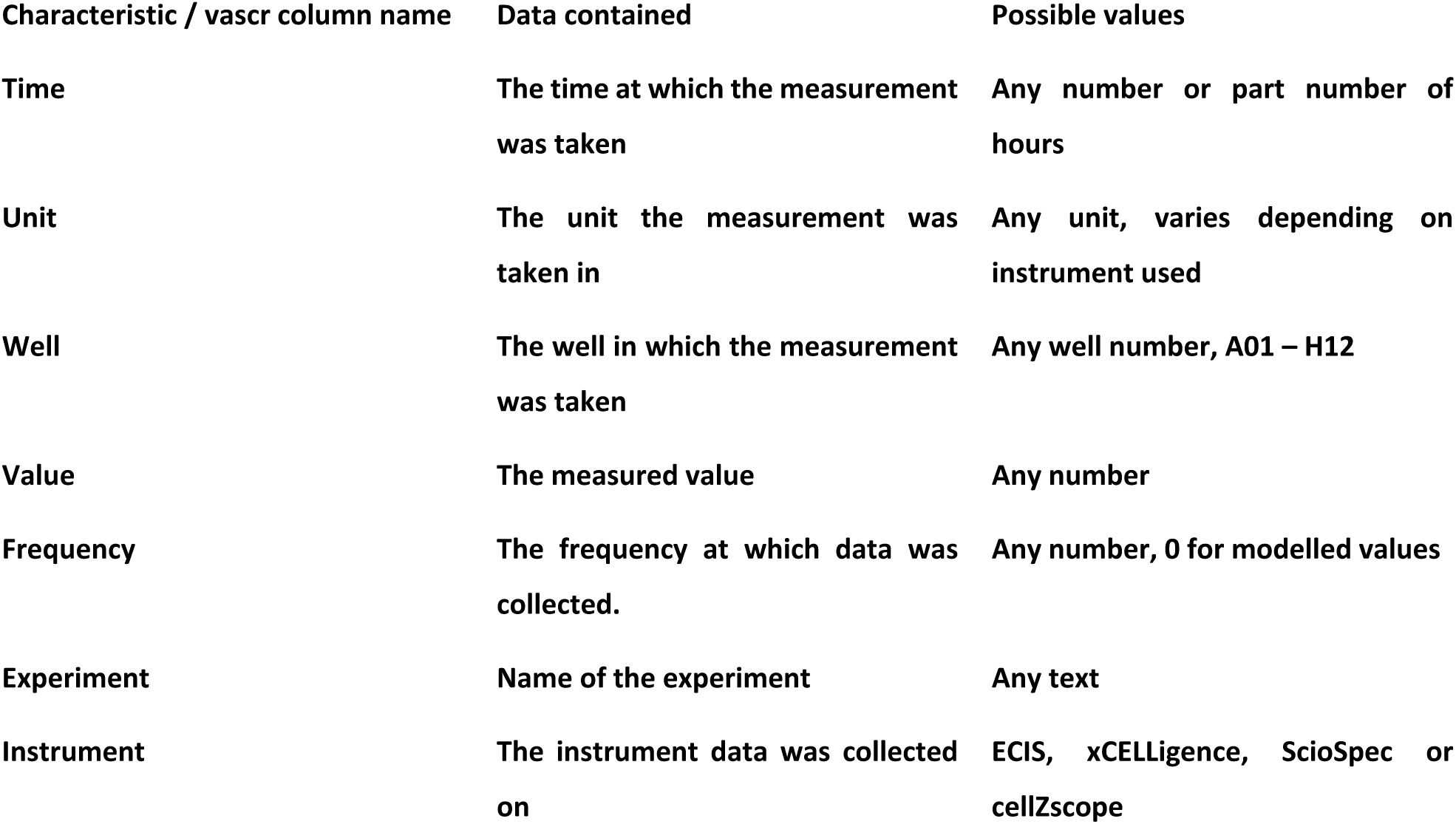
Data types which are imported by vascr.

### 3.2. Resampling provides a consistent time series without compromising data integrity

Examining the raw impedance-sensing data revealed that each well was measured sequentially rather than simultaneously (Figure 1A). Zooming into the first two hours of data collected from hCMVEC cells seeded at 20 000 cells/well revealed that well E01 was sampled slightly before E02. This slight offset in timings was continued across all wells of the plate (data not shown). Whilst these time differences are small, they are a necessary engineering compromise to allow one set of expensive impedance spectroscopy circuitry to measure all wells in the plate. However, the lack of a common measurement time for all wells means that measurements from technical replicate wells cannot be directly averaged. Hence, linear interpolation was applied using the vascr_resample_time function to resample the data. This function uses R’s built in approx function to linearly interpolate between adjacent timepoints, thereby estimating the values which would have been measured if all wells in the plate were measured simultaneously (Figure 2B) [73]. Resampling impedance-sensing data is not new, and is implemented within the Applied Biophysics ECIS software, and is a necessary step before modelling Cm, Rb and α parameters [12][71]. Furthermore, we improved the Applied BioPhysics implementation by removing interpolation when the experiment is paused for the addition of treatments. The change ensures that rapid changes after cellular manipulation, such as that occurring at 47 hours in Figure 1C were captured accurately.

**Figure 1.**
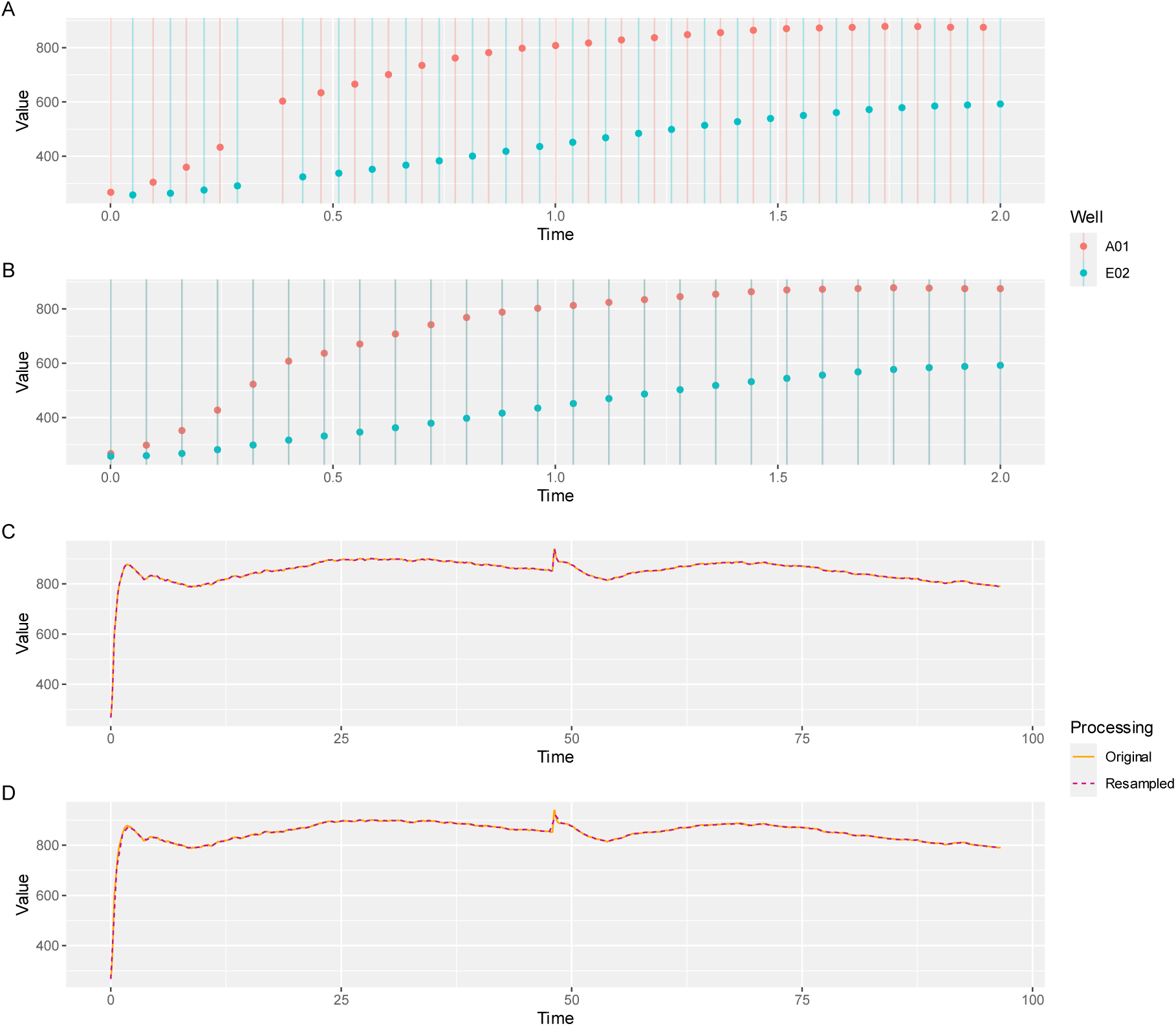
Resampling corrects for small differences in time during the acquisition of data, thus allowing for the direct comparison between technical replicates. hCMVEC cells were seeded at a density of 20,000 cells/well and monitored using ECIS technology for 95 hours. A) Raw data captured from two wells showed a small difference in acquisition time. B) Data from panel A, aligned by resampling at the same rate at which the data were originally acquired. C) Throughout an extended experiment, data resampled at the same acquisition rate maintains an identical profile to the original data. D) Data resampled at a substantially lower rate maintains the overall profile of the original dataset.

**Figure 2.**
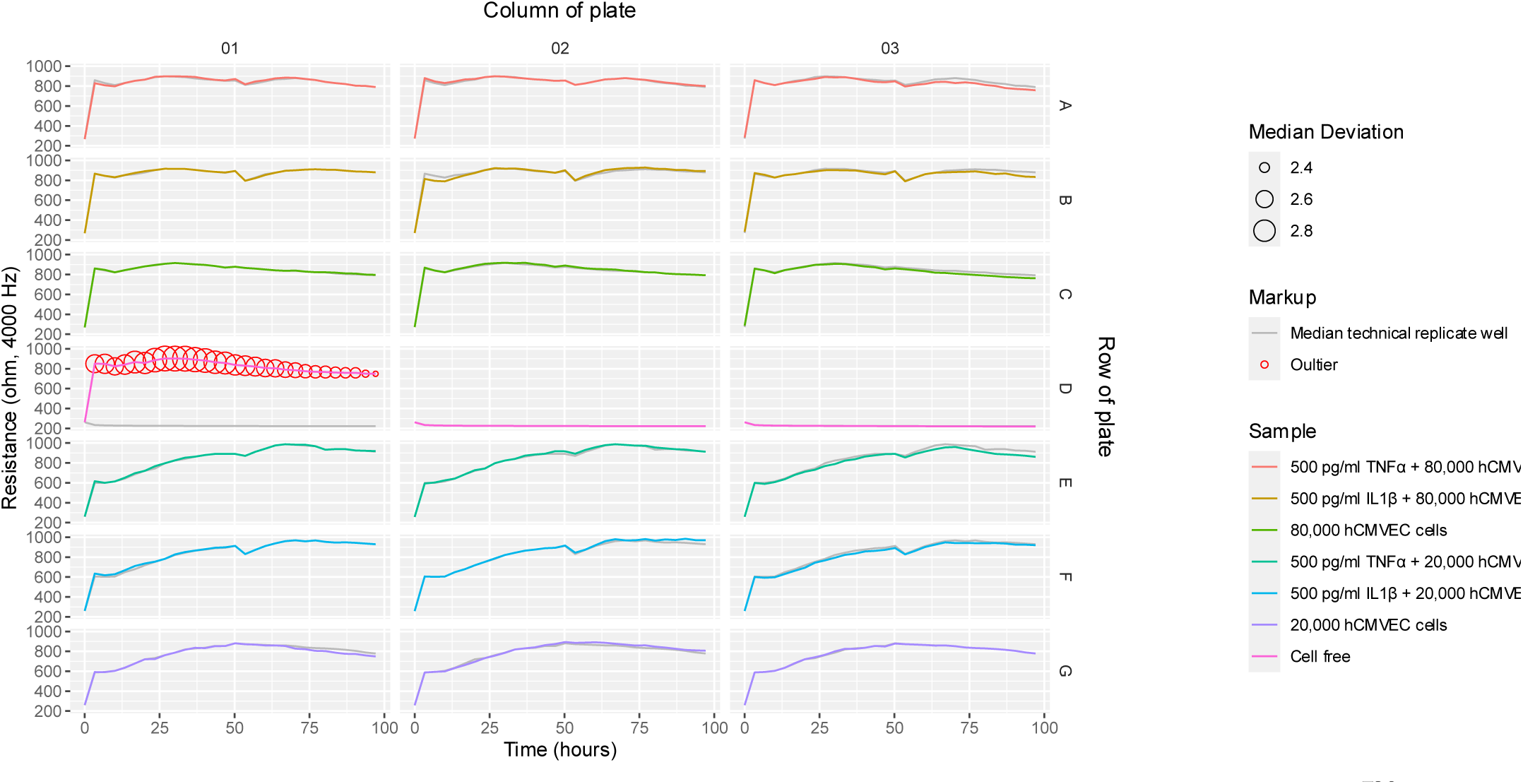
vascr_plot_grid allows for the rapid identification of erroneous data. hCMVEC cells were seeded at various densities and monitored using ECIS technology for 95 hours. After 48 hours, cytokines or paired vehicle controls were introduced. Data shows the profile of each well (coloured), compared to the median well for each treatment condition (grey). Timepoints identified as erroneous are highlighted with red circles. Vascr can plot data combined from multiple technical and experimental replicates, thus highlighting any variation

To confirm the interpolation was accurate, both raw and resampled data were plotted in Figure 1C. Both curves overlaid perfectly, indicating that resampling had no perceptible effect on the underlying data (Figure 1C). Formal statistical testing of the original and resampled values using the vascr_find_resample_frequency function confirmed the curves were identical, with a change in area under the curve of less than 0.01%, R2 of 0.998 and a cross correlation of 0.999. Similar results were observed for all wells and across all four impedance-sensing instruments compatible with vascr (data not shown).

An additional advantage was that resampling could also correct for small differences in measurement time between replicate experiments. These small differences between replicate experiments were common; introduced due to differences between instruments, changes in the number of wells measured and transient connection issues such as the short gap at 0.4 hours observed in Figure 1A. Resampling provided a robust method to correct for these small differences and allowing for consistent data analysis within and between experiments.

Resampling also reduced the computational load of processing impedance sensing data. A 100-hour experiment, such as that in Figure 1C with measurements every 4.3 minutes, generated 1395 datapoints. Whilst this high frequency provides exceptional time resolution to detect rapid changes in cellular behaviour, it substantially slows down data analysis. This limitation can be overcome for fast responses by discarding data collected outside of the short period of interest. However, many biological stimuli occur over a period of several days [6]. As the speed of these responses may not be known in advance and ECIS measurements are non-invasive, it is often desirable to capture these data at the highest possible rate and later resample the data to reduce the number of timepoints and ease analysis. The vascr_find_resample_frequency function can identify an appropriate resampling rate by repeatedly resampling the data at progressively lower rates and comparing the results to the original data. To ensure all characteristics of the underlying data were maintained, three statistical tests were conducted using the stats package [44], namely, cross correlation, change in area under the curve and R2. The rate selected for resampling was the lowest measure where the values from all three statistical tests remained above 0.99, representing an exceptionally good fit. For the data presented in Figure 1D, 151 time points were required. To confirm that the data remained unchanged, the original dataset containing 2,784 timepoints is overlaid with the resampled data containing 151 timepoints in Figure 2D. Whilst the data resampled at a lower rate does not fully capture the high-frequency variations in the cellular response, and implicitly averages out cellular micromotion, it accurately captures the overall shape of the curve [3]. This may be desirable in some situations, as random cellular micromotion may obscure the underlying biological changes [3]. Furthermore, dealing with fewer points is desirable, as it substantially speeds up computational processing and plotting operations. Hence, whilst it must be used with care, resampling provides a good approach to speed up calculations across large experiments with long durations.

### 3.3. Data cleaning objectively removes erroneous data, which occurs due to technical issues

Finally, data needs to be cleaned before it can be further analysed. Cleaning involves the removal of datapoints where there has been a clear technical error, ensuring such values do not distort subsequent analyses [74, 75]. The technical complexity of impedance sensing means that some wells can experience instrumentation errors, such as insecure connections with the array station, scratching of the electrodes or introduction of bubbles by pipette tips during the addition of treatments or inconsistent cell seeding. Whilst good technique can reduce these errors, preventing them entirely is challenging [6, 45]. However, as these technical errors are unrelated to the biological treatments under investigation, they should be removed before further analysis [74].

Instrumentation issues can be rapidly identified by eye using the vascr_plot_table function (Figure 2). vascr will plot the measurements made for each well, with each line colour-coded by the treatment applied. To allow for comparisons between replicate wells with the same treatment, vascr determines which of the replicates is the median value at most timepoints and overlays this well’s trace on the other technical replicates in grey. This display allows the user to visually confirm that the data was fully imported and confirm that the correct treatments were defined for each well, and identify any timepoints or traces with technical issues. Most technical replicates were highly consistent. However, well D01 increased over time, suggesting that it was erroneously seeded with cells rather than remaining cell-free like D02 and D03. It is also clear that data was not collected from well H3, in this case, due to a connection issue between the plate and ECIS array station. Whilst not present in this experiment, we have occasionally detected sudden spikes in resistance with no biological rationale or increased variability in edge wells. These are usually due to connection issues and low incubator humidity, respectively, and should be removed [45, 74, 76].

Whilst much of this erroneous data is obvious from visual inspection of the data, any manual data cleaning process is exposed to explicit or implicit human bias. Furthermore, when various treatments are applied across multiple 96-well plates, manually evaluating the trace from each well becomes time-consuming and error-prone. Therefore, we developed a novel standardised metric. Measures based on standard deviation are inappropriate, as these are strongly affected by extreme outliers, such as the instrumentation issues we are attempting to detect [75]. As a more robust metric of deviation, vascr calculates the difference between each value and the median of the technical replicate wells from the experiment [77]. If this deviation was greater than 34% of the value of the median of the technical replicates at any time point (approximately 1 standard deviation), this sample was deemed to be an outlier and denoted with a red circle (Figure 2 Well D01). We have used this filtering approach extensively and found it to reliably detect outliers, without errantly highlighting data which could reasonably be considered as biological variation [6]. Whilst we have not found it necessary, the 34% cutoff can be tuned for particular applications and then applied uniformly to all subsequent experiments to ensure impartiality. Once identified, outliers can be excluded using the vascr_exclude function in vascr.

Once imported, the values measured in each well over time can be plotted (Figure 3A). This is achieved by summarising the data using the vascr_plot_line function. In Figure 3A, the value of each well throughout the experiment was plotted, with different experimental replicates indicated by different line styles and different experimental treatments indicated by different colours. Whilst this plot highlights the quantity of real-time data collected using ECIS, the sheer volume of data makes interpretation challenging.

**Figure 3.**
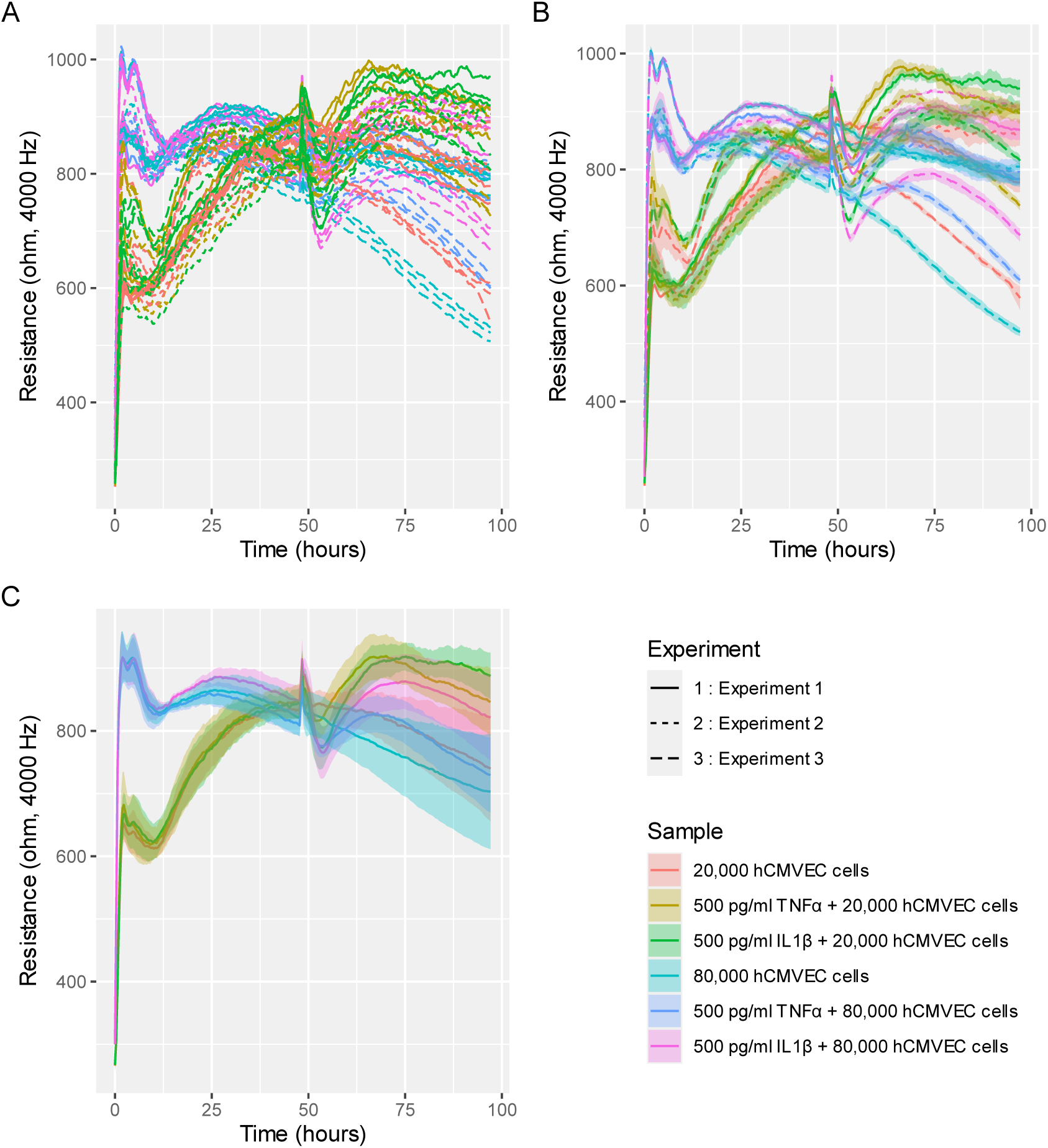
Datasets can be summarised to various levels to best represent the underlying data. hCMVEC cells were seeded at various densities and monitored using ECIS technology for 95 hours. After 48 hours, cytokines or paired vehicle controls were introduced. Data shows the same data presented as A) individual wells, B) average ± SEM of each technical replicate and C) mean +- SEM of three experimental replicates.

To visually simplify these data, vascr can average the technical replicates within each experiment (Figure 3B). This display highlights that the responses to each treatment were similar for each experiment; there were slight differences in the shapes of the curve and absolute values measured between experimental repeats. Whilst this implies that the observed responses were reproducible, the slight differences made the data difficult to interpret as a whole.

To address this challenge, vascr can use the vascr_plot_line function to generate the averages from all replicate experiments (Figure 3C). To calculate these data, the mean of the technical replicates in each experiment was first calculated, and these values were combined to calculate an overall mean ± SEM across the three replicate experiments. This approach is superior to simply averaging all nine technical replicates in a single step, as it reduces the impact of the technical variation and random noise within each experiment on the overall mean, providing a better representation of the biological variation between replicate experiments [75, 78, 79]. Furthermore, averaging only the final data from each experiment conservatively estimates the degrees of freedom, preventing pseudo replication from underestimating the inter-experimental variation [75, 78]. Overall, vascr’s summary level ribbon plots provide a visually straightforward way of visualising the difference in variation within complex experiments.

### 3.4. Normalisation by vascr enables the comparison of treatment responses across multiple experiments

Looking at the mean measurements only, Figure 3C indicates that each cytokine has a substantial effect on brain endothelial microvascular integrity. This finding is consistent with previous literature [6, 23, 35]. However, the substantial variation in the baseline values measured in each well before the addition of cytokines (Figure 3B) results in the large variations in the standard error of the mean (Figure 3C). This variation makes it challenging to interpret the reproducibility and biological relevance of these cytokine responses. Although this variation was reduced as far as practicable by employing experienced personnel and robust protocols, variations in baseline resistance of up to 15% were measured [45]. This level is variation is consistent with previously published findings using ECIS and multiple other commercially available impedance sensors [6]. As impedance sensing is precise, using one set of lock-in amplification electronics to measure all wells, this variation does not stem predominantly from the instrumentation itself [5]. Instead, given that the variation has a random distribution which varies between experiments, it likely stems from random factors unique to each well. These random factors include variation in the seeding and growth patterns of the cells, contact resistance within the connections between the impedance culture ware and array station and manufacturing variation in the culture ware itself. As further controlling this variation within the experiment is not technically feasible [45], a mathematical correction is required [79].

Baseline differences can be mathematically corrected through a process of normalisation [79]. During this process, the value measured at each time point is divided by or subtracted from the value of the same well measured at a key time point. vascr can rapidly carry out these calculations, using the vascr_normalise function. In this experiment, vascr divides the measured value at each time point by the value of the same well at a nominated time point, usually selected to occur just before the treatment is applied. This per-well normalisation is highly effective, as it is able to use impedance sensing’s high temporal resolution to use each well as its own control, elegantly correcting for baseline differences in each well. After normalisation, the average values for each experiment were then calculated. Comparing the normalised results in Figure 4C to the raw data in Figure 4A, the consistency of the responses of the brain microvascular endothelial monolayers to the cytokine treatment is substantially clearer. Figure 4D shows the overall averages of these data, with normalisation carried out before means were calculated using the method described in the previous section. The resulting normalised and averaged data show minimal variation before treatment, as would be expected from identically prepared wells. Whilst there is still substantial error observed towards the end of the experiment, this represents the variation observed between individual replicates in the raw data presented in Figure 3A. Overall, vascr’s process of normalisation and averaging captures the biological variation in a visually simple way.

**Figure 4.**
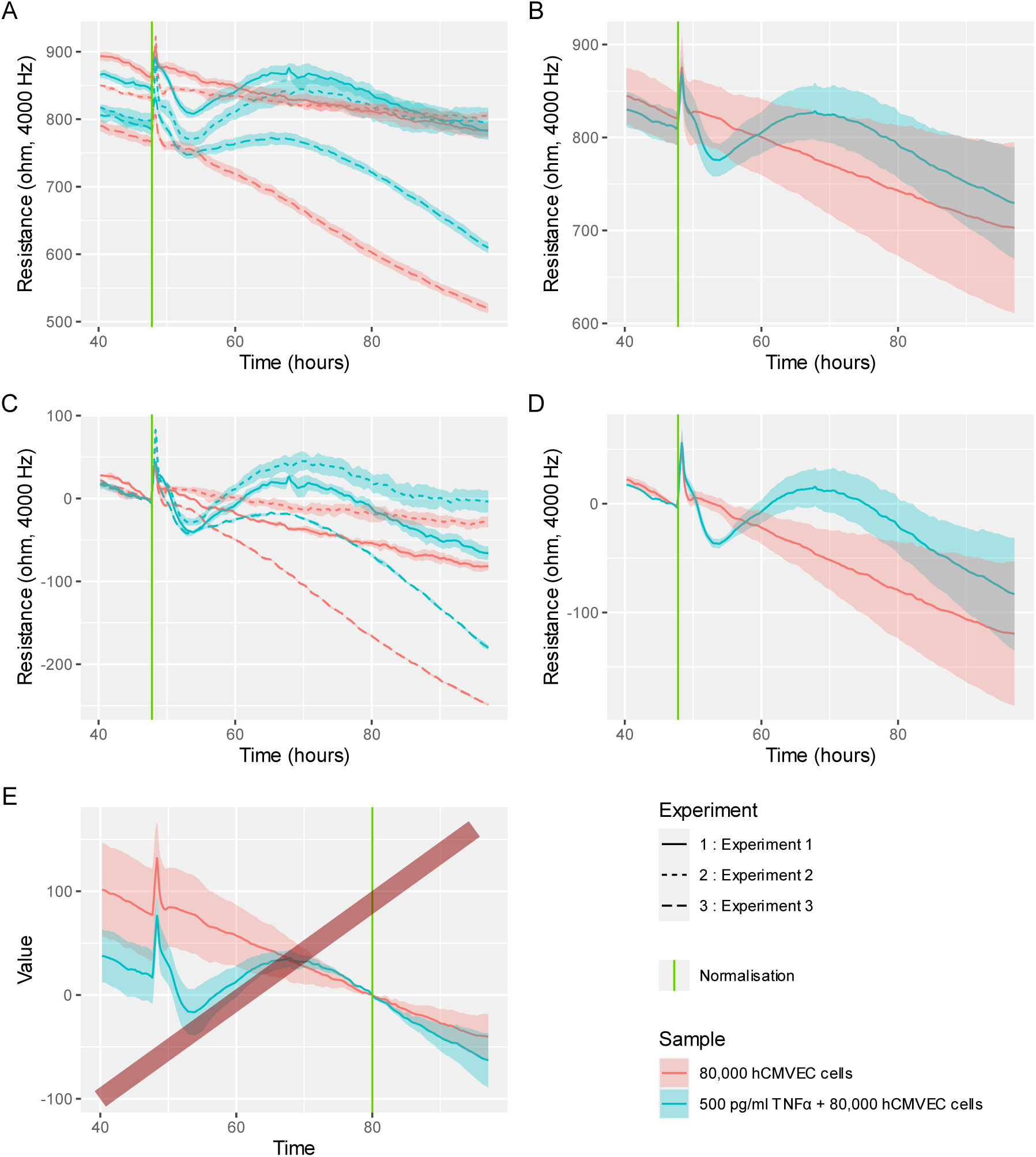
vascr can calculate fold change in resistance from a pre-treatment time point, reducing the baseline difference between replicates and treatments in different experiments, therefore emphasising the difference due to experimental treatment. hCMVECs were seeded at 62,500 cells/cm² and incubated for 48 h until confluent. The cells were then treated with the TNF-α or vehicle and monitored using ECIS for a further 48 hrs. A) The mean ± SEM of three technical replicates for each experimental repeat B) The combined mean ± SEM of all three experiments C) The fold change ± SEM of three technical replicates for each experimental repeat D) The combined fold change ± SEM across all three repeat experiments. E) The same data, presented inappropriately normalised to 80 hours, substantially after treatment is applied. A red bar has been added to this plot to emphasise that this analysis is inappropriate.

Whilst normalisation is a powerful tool for reducing intra-experimental variation, inappropriate usage can result in misleading conclusions. Figure 4E shows the dataset in Figure 4D, however as an example of inappropriate normalisation, data was normalised to the values measured at 80 hours. This time point is inappropriate as 80 hours is well after the application of the cytokine, and therefore, cytokine-treated cells already show substantially reduced barrier integrity. Normalisation artificially removes this reduced integrity, warping the experimental effect. If interpreted in isolation, this data implies that cytokine-treated cells had a substantially lower barrier integrity before cytokines were introduced, and barrier integrity was restored by the cytokine addition. This conclusion is clearly inappropriate as it does not align with the un-normalised data in Figure 4A, previous studies of brain endothelial monolayers [6, 23, 35] and the known biological function of TNFα [35]. To prevent these inappropriate conclusions, normalisation should only be applied to wells which have been treated identically and before any treatments are applied. Secondly, normalisation inherently removes variability. Whilst this may make inferring the effects of a treatment easier by reducing random variation, care must be taken to avoid obscuring biologically important variation such as donor variability or systematic plate effects. Furthermore, as normalisation can make technical errors less obvious, care must be taken to ensure these are removed to prevent artificial inflation of standard error. Therefore, the point at which normalisation is applied should be clearly notated and justified, occur before the addition of treatments, and only be applied to data which has had technical errors removed. By adhering to the guidelines, normalisation can provide a useful tool to highlight the features of the data which represent the genuine biological response.

### 3.5. ANOVA analysis allows robust testing for the significance of differences at distinct timepoints

Whilst plotting the data provides insight into the effect of treatments, formal statistical testing provides an impartial way of demonstrating these hypotheses [75]. Whilst formal testing of Transwell tracer and EVOM assays, which give a readout of barrier integrity at a limited number of time points is common, similar analyses of real-time impedance sensing data are utilised but have not yet become ubiquitous [6, 10, 15, 17, 20, 80, 81]. Whilst some stimuli cause large differences in barrier integrity, which are clearly biologically and statistically significant, formal statistical testing of smaller differences may be required to confirm that the differences observed are genuine [17, 18, 45].

Two-way ANOVA provides a robust way to test if a treatment resulted in a statistically significant difference in the impedance measured [75, 82, 83]. Two-way ANOVA analysis was selected, as it models both variation between experiments and between experimental treatments [75]. Separating these two sources of variation allows two-way ANOVA to correct for the baseline differences observed between repeat experiments outlined in Figure 4, and prevents inter-experimental variation from erroneously inflating the calculated P-value. As normalisation will artificially deflate this inter-experimental variation, ANOVA should only be conducted on raw datasets[84]. To prevent pseudo-replication due to repeated measures in the same experiment, vascr takes a conservative approach to dealing with technical replicates. The vascr_summarise_anova function averages technical replicates on a per-experiment level before ANOVA analysis is run.

Whilst powerful, ANOVA analysis is only robust when four key assumptions are met. Whilst ANOVA is robust, reliably estimating P values even with moderate deviations from ANOVA’s assumptions [78,85], the presence of outliers or inappropriate experimental design can skew these analyses [75, 83, 85]. Therefore, whilst often overlooked, it is important that these assumptions are verified [75, 84]. This analysis can be conducted using either the ANOVA tab of the vascr GUI or using the vascr_summarise_anova function. The output of this function is presented in Figure 5.

**Figure 5.**
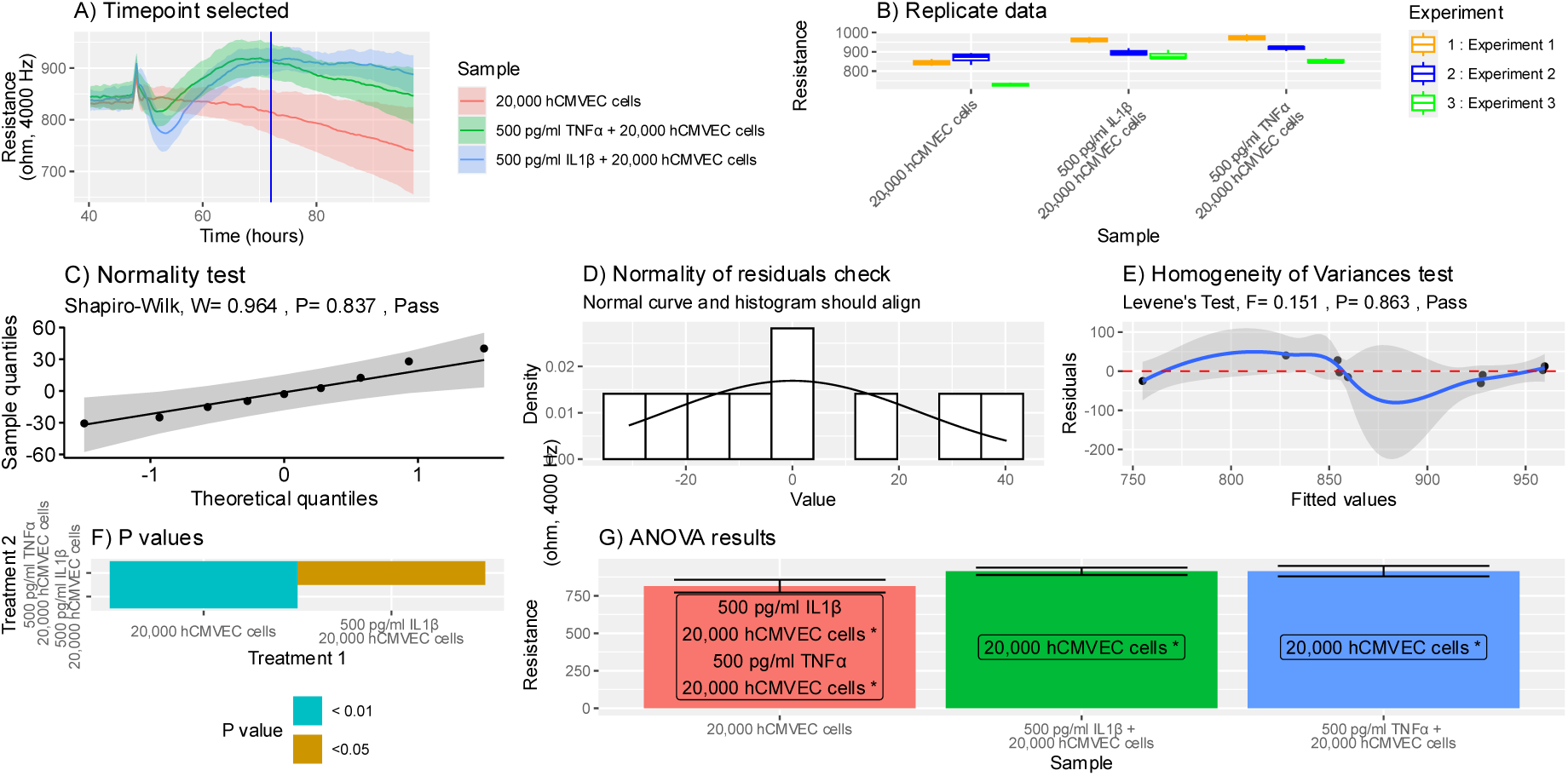
vascr can conduct ANOVA analysis on data at key time points. hCMVEC cells were seeded at a density of 20,000 cells per well and monitored using ECIS technology for 95 hours. After 48 hours, cytokines or paired vehicle controls were introduced. Full ANOVA analysis was then conducted on the experiment. A) Analysis was conducted at 72 hours, as denoted by the blue line. B) Technical variation within each experiment. C) Shapiro Wilks Normality test and Quatntile Quantile plot and D) residuals overlaid with a theoretical normal distribution confirmed normality. E) Levene’s test and a residuals vs fitted plot confirm heteroscedasticity. F) Heatmap of P values and G) bar graph annotated with significance values calculated using Tukey’s HSD test.

ANOVA’s first assumption, that data are independent, is satisfied primarily by experimental design. Independence is satisfied by conducting repeat experiments, each conducted on a separate day, and where possible, utilising separately prepared reagents and cells. Furthermore, standard laboratory procedures should be employed to prevent systematic errors in pipetting and cross-contamination between treatments [45]. To further reduce systematic error, treatments should be applied to randomly assigned wells. As we have observed that edge wells and particular regions of the plate may differ, it is best practice to randomly alter the layout of samples between each repeated experiment [45].

The remaining three ANOVA assumptions, homogeneity of variance, normality of residuals and sphericity, are more challenging to address with experimental design alone. Therefore, these assumptions should be verified mathematically. Whilst often overlooked in impedance sensing assays [75, 82, 83], vascr makes verification trivial by automatically conducting the required calculations and plotting for these tests.

To confirm that the correct data has been selected, vascr_plot_anova first generates a line plot of the underlying data (Figure 5A). This visualisation of the data is critical to reducing human error by ensuring that the intended data, treatments, timepoints and experiments are being fed into the ANOVA analysis[31]. In this example, Figure 5A shows that that the cytokine dataset, previously visualised in Figure 4, has been correctly imported. The blue line indicates that calculations will be performed at the 72-hour timepoint, 24 hours after the addition of cytokines. This point is of interest, as it is when the maximal endothelial response to the cytokines versus the control was observed.

Secondly, ANOVA’s assumption of homogeneity of variances and sphericity was confirmed. To be spherical, the variation within each group should be similar. By inspecting the data in Figure 5B we can see that the ranges of each treatment differed by less than 50%, suggesting variation was similar across the experiment [83]. This homogeneity of variances was then formally tested using the Levene test (Figure 5E). Levene’s test resulted in a P>0.05, meaning we have no evidence that the standard deviations differ between treatment groups [77]. Finally, a fitted vs residuals plot was generated to identify any outliers which might substantially affect the standard deviation and artificially deflate P values (Figure 5C) [85]. As all points were roughly linear and no particular points were substantially further from the mean, we can conclude that no erroneous points are present. This is to be expected, given the removal of technical errors previously discussed.

Vascr next verifies ANOVA’s fourth assumption, normality of residuals was confirmed by running the Shapiro-Wilk test (Figure 5C). In this case, P>0.05, confirming that we do not have any evidence to demonstrate that the data deviates from normal [77]. Normality can also be confirmed visually by a Q-Q plot following a linear distribution (Figure 5C), and a histogram of data roughly following a normal distribution (Figure 5D) [85]. Whilst data should be mostly uniform, the Shapiro-Wilk test becomes highly sensitive to departures from normality with higher sample sizes and may present false positives when many treatment conditions are compared. However, increased samples make ANOVA more robust to violations of this assumption [75, 82]. Therefore, analysis can be continued even if this test fails, if the sample size is relatively large and samples were assigned randomly from a pool of wells prepared identically.

Having determined that the ANOVA’s assumptions are met, we can draw conclusions from the results of the ANOVA test. Figure 5F shows a heatmap, showing that the vehicle is highly significant (P<0.001) versus both cytokine treatments. However, at this time point, the two cytokine treatments were not significantly different from each other. Finally, the data and statistical testing is presented in a straightforward bar plot as Figure 5 D. This plot is a straightforward representation of the data, and is annotated with which samples are statistically significant versus each other. This plot is also important, as it shows the absolute size of any changes detected. This is important information, as even though a small difference may be highly reproducible and therefore statistically significant, it may not reach biological significance. In this case, both cytokines are known to have clinically significant phenotypes by causing substantial changes in a wide range of cellular functions, including substantial changes in endothelial integrity [35, 86, 87]. Therefore, this change is both biologically and statistically significant. To simplify interpretation, only two cytokine treatments are shown in Figure 5. However, this ANOVA approach can be scaled up to account for large numbers of simultaneous treatments [6, 11, 16, 20–22].

Whilst all treatments applied can be compared, it is often more sensible to compare each sample with an untreated control. In the cytokines example, it is more rational to ask whether each cytokine influences the endothelium, rather than which of the cytokine responses is significantly different from the others. This distinction is particularly important when screening drugs or cytokine responses, where the efficacy of an agent is substantially more important than each agent’s effect relative to the others. Fewer comparisons are simpler to interpret, whilst statistical power is increased due to the reduced number of comparisons required [88, 89]. This pairwise analysis can be carried out by specifying a reference in vascr. vascr will carry out an identical ANOVA analysis; however, Tukey’s HSD is substituted for the Dunnett’s test without correction using the glht function at each time point[90]. Multiple time points can then be combined, and multiple comparisons adjusted for using the Bonferroni correction[89]. vascr will then superimpose these results onto a normalised line graph, showing the significance of p-values versus the control. The control itself is also clearly marked with a + symbol (Figure 6). Whilst the data is normalised to reduce the effect of technical variation, as previously discussed, the statistical testing is performed only on the raw data. In Figure 6, statistical testing was conducted 5 and 22 hours after the addition of cytokines, timepoints selected to capture the largest differences in barrier integrity compared to the control. Overall, this display provided an elegant solution, combining formal statistical testing for a difference in magnitude without losing the real-time context and clarity of normalised data unique to impedance sensing.

**Figure 6.**
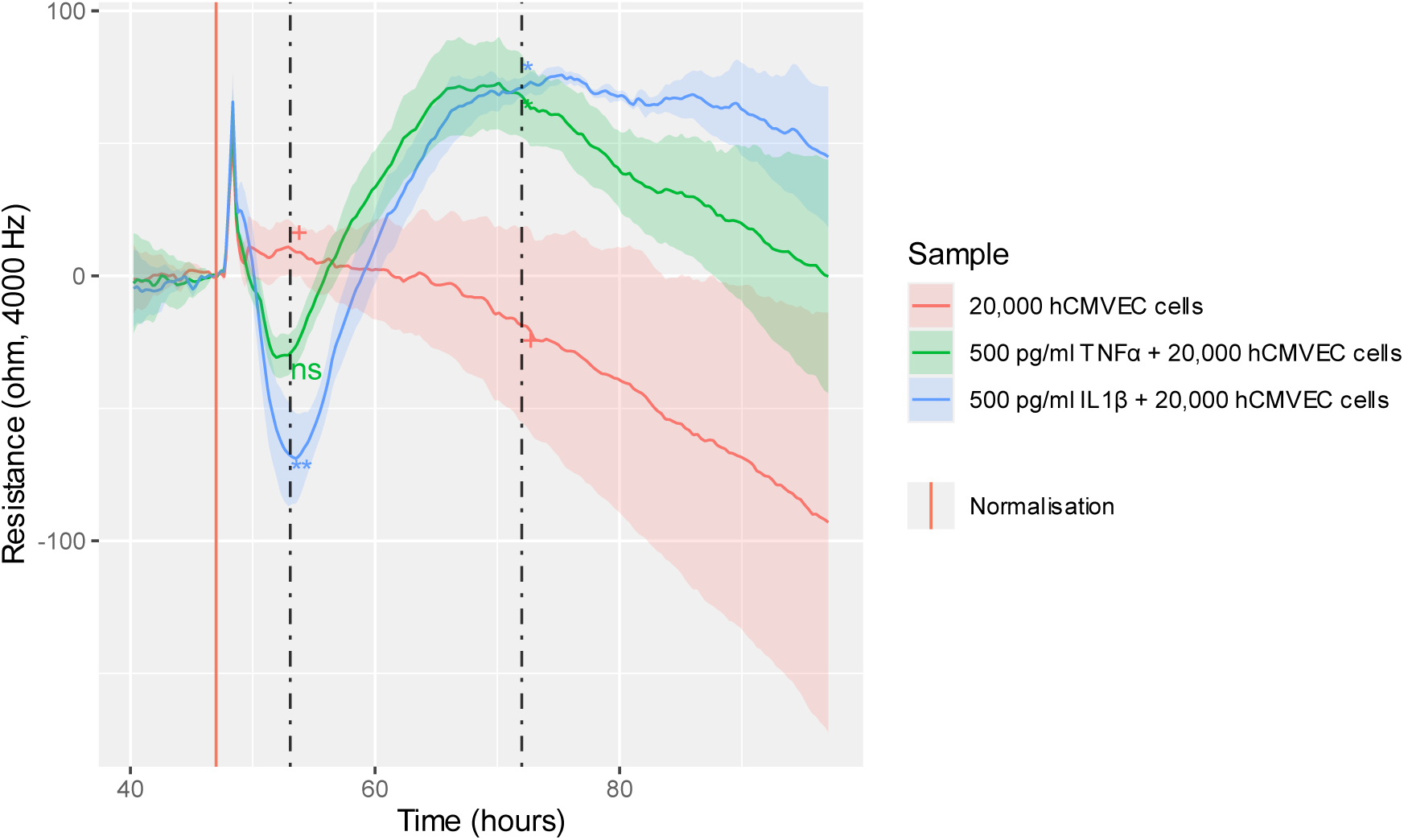
Statistical testing comparing various treatments can be directly applied to the plot at key time points. hCMVEC cells were seeded at a density of 20,000 cells per well and monitored using ECIS technology for 95 hours. After 48 hours, cytokines or paired vehicle controls were introduced. Full ANOVA analysis was then conducted on the experiment. Dunnett’s test with Bonferroni’s correction was carried out 5 and 24 hours after the addition of cytokine (* p < 0.05, ** p < 0.01, + reference value).

### 3.6. Cross-correlation analysis allows for comparisons of the overall shape of the curves

Cross-correlation provides a powerful way to compare the shapes of curves. Whilst ANOVA allows for absolute differences to be compared, it is often more informative to compare the shapes of the underlying curves. Cross-correlation condenses the difference between the shapes of the two-time series into a single value [42]. This can be seen in Figure 7A, which shows a set of test curves and their corresponding cross-correlation values. The black reference curve is identical in shape to the red curve but has a larger magnitude. Hence, the shapes are the same, and the cross-correlation is 1. The green curve is the exact inverse of the black reference curve, therefore they have a cross-correlation of −1. Finally, the blue curve has a substantially different shape, and a cross correlation of −0.227, indicating that it is mostly unrelated but slightly inverted versus the black control.

**Figure 7.**
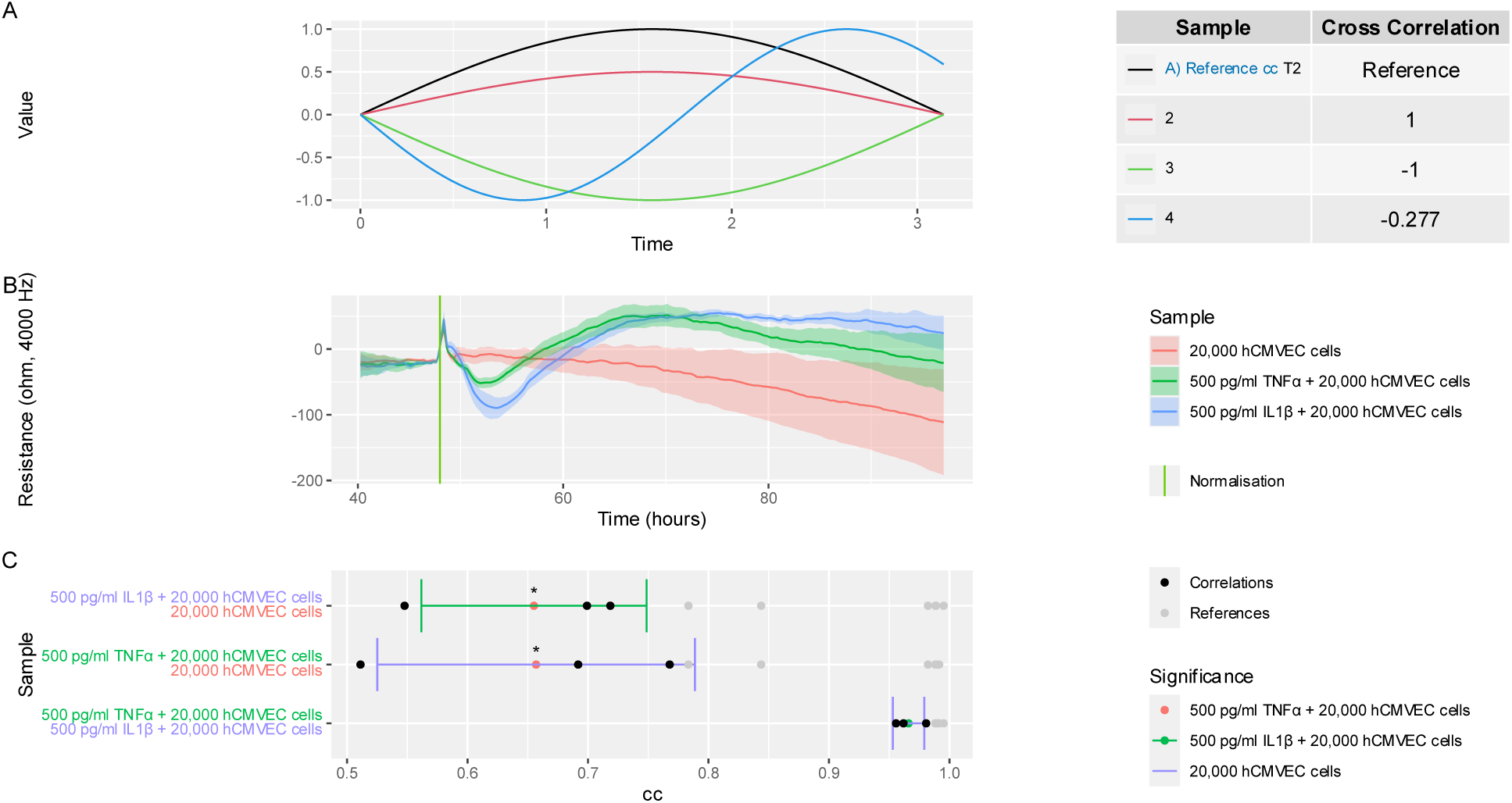
Cross-correlation allows for the shapes of various curves to be compared. A) Example data, referencing an example dataset showing that curve 2 (data with an identical shape but different magnitude) and curve 1 have a cross correlation of 1, whereas curve 3 (which has an opposite shape) and curve 1 have a correlation of −1. Curve 3 (which has a somewhat different shape) has an intermediate cross-correlation when compared to curve 1. B) Changes in Rb due to treatment with cytokines. C) Cross correlation shows the mean +- SEM of the cross correlations calculated between each curve (black dots). B) hCMVEC cells were seeded at a density of 20,000 cells per well and monitored using ECIS technology for 95 hours. After 48 hours, cytokines or paired vehicle controls were introduced. Ribbon plots show the mean ± SEM of three independent experiments, each of which was conducted in triplicate. Each curve was also compared to other wells which received the same treatment (grey dots). C) Cross correlation results and statistical analysis using Welch’s T test against an autocorrelation control (* p < 0.05). All results are derived from technical triplicate values from each of three independent experiments

This cross-correlation approach was then applied to compare the shapes of the two cytokine responses (Figure 7C). Unsurprisingly, the two cytokine responses were the most similar to each other, with a cross-correlation approaching 1. Both cytokines were substantially different from the control, with the TNFα response being more variable than that of IL1β and resulting in a slightly smaller cross-correlation. These observations aligns with the known biology of this process, and our previous analyses of this dataset in the context of comparing impedance sensors [6, 35]. However, the ability of vascr to conduct these analyses automatically allows such analyses to be readily extended across large numbers of samples.[6, 35]

Whilst cross-correlations alone provide insight into how similar responses are, it may also be insightful to run formal statistical testing of the significance of any differences in cross-correlation detected. To do this, as with previous analyses, technical replicates in each experiment were averaged. Cross-correlation was then conducted between each treatment and each treatment in the other technical replicates (Figure 7C, black dots). To give a baseline as to the expected variation in the reference treatment, cross-correlation between each treatment and the same treatment across each experimental repeat was calculated (Figure 7C, grey dots). This correlation between identical treatments across experimental repeats provides a baseline for the level of variation which would be expected if the differing treatments had no effect. These between-treatment and same-treatment values were then compared using Welch’s t-test [77]. Given that the cross-correlation between treatments and within treatments did not have similar variations, Welch’s T-test was used [91]. Multiple comparisons were then corrected for using an fdr approach[89]. The results of this analysis are shown as stars on Figure 7C, which determined that both cytokine treatments were statistically significant versus the control, but not versus each other. Given the ubiquity of comparing the shapes of responses, this analysis has applications in a wide variety of experiments, which may include responses of differing shapes, including drug screens, cancer growth profiles and endothelial responses to stimuli.

## 4. Conclusions

Overall, vascr provides a rapid and effective way to analyse impedance-sensing data. This reduction in manual transposing time increases laboratory productivity, whilst the ability to conduct analysis allows data analysis to be conducted on provisional data, which can then be rapidly repeated by simply re-running the same code on replicate datasets as they are generated. The high reproducibility also reduces the risks of human errors in data processing, such as unconscious bias and data loss. Furthermore, code can be easily audited, allowing for verification of experimental findings by laboratory heads and external reviewers. Finally, as the process is consistent across multiple impedance sensing platforms, users can utilise multiple instruments to select the one best suited to their particular experimental questions. Hence, vascr is already used routinely within our own laboratory, and has wide applicability to impedance-sensing experiments in a variety of fields.

## Author Contributions

Conceptualization: J.J.W.H., C.E.A. and E.S.G.; methodology: J.J.W.H., C.P.U., C.E.A., and E.S.G.; software: J.J.W.H.; formal analysis: J.J.W.H., C.P.U., C.E.A., and E.S.G.; resources: J.J.W.H., C.E.A., and E.S.G.; data curation: J.J.W.H., C.P.U., and E.S.G.; writing—original draft preparation: J.J.W.H.,C.E.A. and E.S.G.; writing—review and editing: J.J.W.H., C.P.U., C.E.A., and E.S.G.; supervision: C.E.A. and E.S.G.; project administration: J.J.W.H., C.E.A. and E.S.G.; funding acquisition: J.J.W.H., C.E.A. and E.S.G. All authors have read and agreed to the published version of the manuscript.

## Data Availability Statement

All code used in this paper is available online at vascr.huc.nz and via CRAN. Raw data files are available from the authors on reasonable request.

## Acknowledgments

We thank the University of Auckland Statistical Consulting Centre for their input and verification of the statistical analysis conducted in this paper. We also thank Jo Dodd for her generous guidance and technical support.

